# Improved Centromere Assemblies for RefGen_v4

**DOI:** 10.1101/2022.03.30.486462

**Authors:** Daniel J. Laspisa, Kevin L. Schneider, Gernot G. Presting

## Abstract

Genome assemblies based on long read sequencing technology have revolutionized the assembly of repeat-rich centromere regions. However, because maize centromeres are highly enriched for the tandem repeat CentC and centromeric retrotransposons (CR), automated genome assembly left gaps even in the excellent **B73 RefGen_v4** reference genome constructed from long-read data. Manual editing of >140 Mb spanning the ten centromeres of maize inbred B73 resulted in the closure of 127 sequence gaps and the addition of >8.4 Mb of previously unanchored sequence (unitigs and reads) containing 24 genes, 2 Mb of CR repeat and 887 kb of CentC without including any additional sequence data. The functional centromeres of five maize chromosomes were closed completely, including a 7 Mb region spanning the extremely CR2-rich CEN2. This improved assembly, **B73 RefGen_v4CEN**, was completed in February 2019 and has been available at https://doi.org/10.25739/7y1p-5169, both as pseudomolecules and as centromere assemblies alone. Thus, the manual editing of existing sequence data significantly improved the centromere regions of the **B73 RefGen_v4** reference genome. These data were used for centromere analyses until the release of RefGen_v5.

## Introduction

Sequencing technology has continued to improve greatly over the past two decades. In maize, the utility of early “long”-read PacBio technology was first demonstrated with the assembly of centromere 10 (using BAC DNA) that resulted in a 1.85 Mb assembly, with a substantial reduction in gap number from 140 to three relative to RefGen_v3, and addition of 50 CRs (Wolfgruber, Nakashima et al. 2016). Until the recent release of the B73-Ab10 (Liu, Seetharam et al. 2020) and B73 RefGen_v5 assemblies (Hufford, Seetharam et al. 2021), RefGen_v4 (Jiao, Peluso et al. 2017) was the highest quality *Zea mays* assembly available, with notably better centromere assemblies than previous B73 reference genomes. However, RefGen_v4 was still fragmented and incomplete in the centromere regions. Manual editing using existing project data was used to close all gaps except those flanked by >10 kb of tandem repeat (primarily the centromere-specific CentC). For each chromosome, we edited the centromere proper, including the ancestral centromere location, all regions on which a neocentromere formed in one or more NAM parents, and the non-recombinant regions consisting of a single or two haplotypes (Schneider, Xie *et al*. 2016).

Frequent neocentromere formation was recently observed in domesticated maize, resulting in the replacement of ancestral CentC-rich centromeres by neocentromeres that are enriched for centromere targeting retrotransposons (Schneider, Xie *et al*. 2016). Although the focus of this project was assembly of the ten functional centromeres of inbred B73 (approximately 20 Mb) as defined epigenetically by the centromere-specific histone H3 variant cenH3, we extended our efforts to include the ancestral centromere locations, as well as the previously pericentromeric regions that had given rise to neocentromeres in one or more of approximately 20 maize inbreds following domestication. These regions had been previously identified via cenH3 ChIP-seq in inbreds B73, Mo17, W64A and most of the NAM lines (Schneider, Xie *et al*. 2016). Moreover, the centromere regions of domesticated maize had previously been found to contain very low levels of genetic diversity, e.g. seven of the ten maize centromeres are represented by only one or two pre-domestication haplotypes in these, otherwise genetically diverse, inbred lines. The edited regions of these chromosomes were extended to the edges of the one (CEN1, CEN4 and CEN6) or two haplotype (CEN2, CEN3, CEN8 and CEN9) centromere regions described previously (Schneider, Xie *et al*. 2016), which resulted in a total of >140 Mb of the maize genome being subjected to manual editing.

## Materials and Methods

### Extraction and annotation of RefGen_v4 centromere and pericentromere regions

MUMmer (Kurtz, Phillippy *et al*. 2004) was used to identify the centromere and major haplotype regions (Schneider, Xie et al. 2016) in RefGen_v4 as defined in RefGen_v2 (Schnable, Ware et al. 2009). Following extraction of these sequences from RefGen_v4 the regions were analyzed using JunctionViewer software (JV) (Wolfgruber and Presting 2010) to visually identify gaps, tandem CentC repeats (Ananiev, Phillips *et al*. 1998), CR elements belonging to subfamilies CR1 through CR6 (Sharma and Presting 2008, Sharma and Presting 2014), genes, direct and inverted repeats, organellar genome fragments, and other retrotransposon junctions. RefGen_v4 pseudomolecules were obtained from NCBI (PRJNA10769), unitigs (provided by the Ware laboratory) and reads (SRP067440) were provided by the Maize B73, AGPv4 Consortium. A subset of the RefGen_v4 sequence data was used to assemble RefGen_v4CEN (**Table 1**) including components of 3,303 high quality unitigs, 145,880 low quality unitigs and 4,047,379 reads.

**Table 1.** Statistics for the subset of RefGen_v4 data used to assemble RefGen_v4CEN. All values represent nucleotides unless noted otherwise.

| Sequence Type | Number of Sequences | Max # of reads per contig | Min # of reads per contig | Median length | Total bases |
| --- | --- | --- | --- | --- | --- |
| High quality unitigs | 3,303 | 11,139 | 3 | 432,132 | 2,099,336,497 |
| Repeat unitigs | 14,338 | 1,587 | 2 | 21,816 | 329,013,878 |
| Singleton unitigs | 131,542 | 1 | 1 | 9,461 | 1,467,086,700 |
| Reads | 4,047,379 | N/A | N/A | 8,630 | 40,351,155,710 |

### Anchoring all assembled unitigs that contain centromere repeat sequence

All high quality unitigs were examined for CentC, CR1 and CR2. Thirty-eight of the 57 CentC-containing high quality unitigs had been placed in the original RefGen_v4 assembly. Thirteen additional CentC-containing HQ unitigs were anchored during the editing process. Three of these were chimeric (utg3669, utg15323 and utg8675) and required trimming of the misplaced sequence (based on RefGen_v2) prior to incorporation into RefGen_v4CEN. Four CentC-containing high quality unitigs, which are comprised of 40% to >80% CentC, remain unplaced because they did not contain unique junctions that could be mapped to a centromere. Two high quality unitigs containing CR2 remain unanchored: chimeric utg115865 (695kb) contains a single CR2 that had been improperly joined at one end and utg129473 (345kb) contains one CR2 near a sequence similar to rDNA.

### Gap Closure

Using the centromere JV images as a guide, unique junctions flanking the gaps in the RefGen_v4 assembly were used to identify overlap, first with singleton or repeat unitigs, then raw long reads or, if needed, using sequence from RefGen_v2. The highest quality sequence containing the junction was imaged with JV and compared visually to the anchored sequence to confirm overlap. A gap was considered closed if a unitig or read covered two unique junctions on either side of the gap. Chimeric unitigs, for which different regions mapped to more than one chromosome or two distinct regions on the same chromosome of RefGen_v2, were split at the chimeric point prior to assembly.

Gaps within the tandem repeats CentC and Cent4 (Page, Wanous *et al*. 2001) were filled by searching for all unique junctions from the corresponding centromeric regions of B73 RefGen_v2. If a CentC or Cent4 containing fragment was flanked by approximately 10kb of CentC, Cent4, or organellar sequence, a gap of 100 Ns was inserted between the fragments.

### Assembly

After all unitigs that could be used to close the gaps were identified, they were imported into Geneious 6.0 (Kearse, Moir *et al*. 2012). The B73 RefGen_v4 assembly region was extracted and broken into contigs at gaps. Gapped regions were assembled using the “Map to Reference” function, aligning one unitig at a time to walk across each gap. The sensitivity parameters varied from unitig to unitig, but “medium sensitivity” was used to assemble most unitigs.

Sequences were manually edited to introduce the least number of ambiguities. Low-quality sequence that overlapped with high quality sequence was trimmed to retain the latter. For chromosomes 2, 4, 5 and 10, overlapping low quality unitigs were trimmed to remove overlap. For chromosomes 1, 3, 6, 7, 8 and 9 unitig overlaps were manually edited for continuity at the nucleotide level. Raw PacBio reads were trimmed if overlapping a unitig or another read to introduce as little error from a raw read as possible. A final JV annotation of the region was constructed for manual verification of the updated assemblies.

## Results and Discussion

The RefGen_v4CEN assembly contains an additional ∼8.4 Mb of sequence data resulting in the closure of 127 gaps (**Table 2**). Approximately 2.9 Mb of centromeric repeat, including ∼1.18Mb of CR2, were added to the regions. In the functional B73 centromeres, 16 gaps were closed and ∼2Mb of N’s were replaced with sequence that included more than 1Mb of CR2 (**Table 3**). Five of the ten B73 active centromeres (CENs 2, 4, 5, 8, and 9) are now complete and contain no gaps. Approximately 5.3 Mb of sequence is estimated to be missing, mostly from CEN1, CEN6 and CEN7. Thus, we show that manual editing can significantly improve assemblies generated from early long read sequencing data.

**Table 2.** Assembly statistics of the centromeres and flanking regions. In total, 127 gaps were closed and ∼8.4Mb of sequence (almost half of which replaced Ns) was added to RefGen_v4CEN. All values represent nucleotides unless noted otherwise.

| CEN | Manually Edited Region (Mb) | # Gaps Closed | Change in Length (v4CEN – v4) | Replaced "N" | CR1 Added | CR2 Added | CentC Added |
| --- | --- | --- | --- | --- | --- | --- | --- |
| CEN01 | 13.949 | 15 | 171,109 | 458,285 | 73,294 | 37,834 | 112,428 |
| CEN02 | 7.001 | 9 | 9,566 | 683,782 | 52,790 | 221,512 | 14,503 |
| CEN03 | 9.998 | 9 | 710,338 | 443,926 | 150,893 | 218,558 | 179,428 |
| CEN04 | 20.3 | 29 | 626,614 | 423,723 | 145,883 | 237,414 | 0 |
| CEN05 | 25.839 | 10 | -22,609 | 773,775 | 30,175 | 81,505 | 48,302 |
| CEN06 | 14.29 | 4 | 511,716 | 6,539 | 180,632 | 93,497 | 122,457 |
| CEN07 | 14.238 | 14 | -72,634 | 355,942 | 39,372 | 52,513 | 94,547 |
| CEN08 | 10.353 | 11 | 279,166 | 401,666 | 50,736 | 164,266 | 99,367 |
| CEN09 | 11.899 | 11 | -50,855 | 113,232 | 50,628 | 6,125 | 100,027 |
| CEN10 | 12.929 | 15 | 2,274,592 | 393,291 | 120,038 | 69,483 | 116,841 |
| **Total** | **140.796** | **127** | **4,437,003** | **4,054,161** | **894,441** | **1,182,707** | **887,900** |

**Table 3.** Assembly statistics of the functional centromeres of B73. N’s were added to CENs 1, 6 and 7 in tandem repeat gaps of unknown length resulting in an increase in total ambiguities in the major haplotype regions of CEN1, 6 and 7. Loss of CR2 sequence in CEN9 was due to removal of chimeric CR2 sequence. All values represent nucleotides unless noted otherwise.

| CEN | SIZE (Mb) | Missing Sequence (Mb) | # of Gaps | Replaced "N" | CR1 Added | CR2 Added | CentC Added |
| --- | --- | --- | --- | --- | --- | --- | --- |
| CEN01 | 0.27 | 1.52 | 3 | -1,885 | 19,332 | 36,995 | 67,206 |
| CEN02 | 2.11 | complete | 0 | 683,682 | 52,100 | 215,591 | 14,537 |
| CEN03 | 1.56 | 0.17 | 4 | 372,341 | 111,812 | 186,115 | 104,141 |
| CEN04 | 1.75 | complete | 0 | 313,321 | 115,093 | 230,098 | 0 |
| CEN05 | 1.96 | complete | 0 | 277,105 | -1,312 | 54,564 | 0 |
| CEN06 | 0.52 | 0.71 | 4 | -1,203 | 104,417 | 48,975 | 90,980 |
| CEN07 | 0.19 | 2.3 | 3 | -1,251 | 27,789 | 37,098 | 53,688 |
| CEN08 | 2.02 | complete | 0 | 357,416 | 14,026 | 145,603 | 0 |
| CEN09 | 1.72 | complete | 0 | 1,004 | 0 | -8,741 | 0 |
| CEN10 | 1.81 | 0.6 | 2 | 70,840 | 102,337 | 63,708 | 57,257 |
| **Total** | **13.91** | **5.3** | **16** | **2,071,370** | **545,594** | **1,010,006** | **387,809** |

Chimeric assemblies are particularly common in centromere regions due to the high density of repeat elements, which increases the likelihood of mis-assembly in regions of relatively low sequence coverage, particularly at the termini of unitigs. In these locations, sequence reads originating from different retrotransposons or tandem repeat sequences can be falsely joined and terminate the contig. These chimeras are identified through analysis of retrotransposon junctions or by cross reference with RefGen_v2 data. Chimeras can exceed 10kb in length and, until the advent of extremely long reads, were one of the major obstacles in assembling high quality centromeric sequence. CR-rich regions containing many nested elements are particularly vulnerable to forming chimeric unitigs, making gap closure difficult.

Centromeric tandem repeats (CentC) remain an obstacle for closure of the active B73 centromeres. Previous karyotyping of B73 revealed CEN1, CEN6 and CEN7 to contain large amounts of CentC (Albert, Gao et al. 2010). These CentC-rich centromeres are difficult to assemble as they lack unique junction markers spaced at intervals short enough to span a single read but may be filled in at a future date when large datasets of even longer reads allow the spanning of large tandem repeat regions.

## Acknowledgements

We thank Doreen Ware and Cold Spring Harbor Laboratory for sharing RefGen_v4 pseudomolecules and unincorporated unitigs and reads prior to publication. The technical support team and advanced computing resources from University of Hawaii Information Technology Services – Cyberinfrastructure, funded in part by the National Science Foundation MRI award #1920304, are gratefully acknowledged. This work was supported with funding from the National Science Foundation grant #1444624, the USDA National Institute of Food and Agriculture, Hatch Multi-state project #1014732 (HAW05037-R), and the University of Hawaii.

